# NFATC4 Promotes Quiescence and Chemotherapy Resistance in Ovarian Cancer

**DOI:** 10.1101/825497

**Authors:** Alexander J. Cole, Mangala Iyengar, Patrick O’Hayer, Daniel Chan, Greg Delgoffe, Katherine M. Aird, Euisik Yoon, Shoumei Bai, Ronald J. Buckanovich

## Abstract

Development of chemotherapy resistance is a major problem in ovarian cancer. One understudied mechanism of chemoresistance is the induction of quiescence, a reversible non-proliferative state. Unfortunately, little is known about regulators of quiescence. Here we identify the master transcription factor NFATC4 as a regulator of quiescence in ovarian cancer. NFATC4 is enriched in ovarian cancer stem-like cells (CSC) and correlates with decreased proliferation and poor prognosis. Treatment of cancer cells with cisplatin results in NFATC4 nuclear translocation and activation of NFATC4 pathway, while inhibition of the pathway increased chemotherapy response. Induction of NFATC4 activity results in a marked decrease in proliferation, G0 cell cycle arrest and chemotherapy resistance, both *in vitro* and *in vivo*. Finally, NFATC4 drives a quiescent phenotype in part via downregulation of MYC. Together these data identify that NFATC4 as a driver of quiescence and a potential new target to combat chemoresistance in ovarian cancer.

## Introduction

Every year approximately 240,000 women are diagnosed with ovarian cancer worldwide, and 140,200 succumb to the disease ^1^. Among all cancers in developed countries, ovarian cancer has the third highest incidence:mortality ratio. Although initial response rates to cytoreductive surgery and primary chemotherapy can be as high as 70%, the vast majority of patients experience a cancer relapse, develop chemotherapy-resistant disease, and die of their cancer ^2^. Consequently, identifying and understanding mechanisms of chemotherapy resistance in ovarian cancer are essential for the development of new therapeutics to prevent relapse and improve overall survival.

Quiescence is defined as a reversible non-dividing state in which cells arrest in the G0 phase of the cell cycle. Adult stem cells are typically maintained in G0 until stimulated to enter the cell cycle and proliferate ^3^. As chemotherapy targets rapidly dividing cells, quiescent stem cells are innately resistant to these therapies ^4^. A striking example of this mechanism can be observed in the hair follicle, where the Nuclear Factor of Activated T-cells (NFAT) family member, NFAT1, drives Cyclin Dependent Kinase 4 (CDK4) downregulation in the stem cell pool to induce a quiescent state ^5^. During chemotherapy, the rapidly dividing follicular cells die resulting in hair loss; however, due to the NFAT1 induced quiescence, the stem cells survive allowing hair re-growth following cessation of therapy.

The role of quiescence in cancer is new area of research. Quiescent cancer stem-like cells (CSC) have been reported in leukemia, medulloblastoma and colon cancers ^6–8^. In some cases, the quiescence is niche dependent, driving CSCs resistance to chemotherapeutics and tumor recurrence ^6–8^. Consequently, successful targeting of quiescent CSCs may be essential for improving cancer cure rates. To date, little is known about regulators of quiescence in ovarian cancer.

We previously reported the identification of ovarian CSC populations defined by the expression of the stem cell makers aldehyde dehydrogenase (ALDH) and CD133 ^9^. Meeting the definition for CSCs proposed by Weinberg and colleagues ^10^, these cells had enhanced tumor initiation capacity, and the ability to both self-renew and asymmetrically divide ^9,11^. In addition, these cells exhibit increased resistance to chemotherapy ^9^. Here, we demonstrate that the NFAT family member NFATC4 (coding for the NFATC4 protein) is upregulated in ovarian CSCs and in response to chemotherapy undergoes cytoplasm to nuclear translocation and with subsequent activation of known NGATC4 target genes. Using two constitutively active NFATC4 constructs, we demonstrate that NFATC4 drives the induction of a quiescent state characterized by (i) decreased proliferation rates, (ii) smaller cell size, and (iii) and arrest of cells in G0 ^12^. Furthermore, induction of NFATC4 conveyed growth arrest and chemoresistance both *in vitro* and *in vivo*. Suggesting, that NFATC4 driven quiescence is in part related to suppressed MYC activity, activation of NFATC4 results in suppression MYC expression, and overexpression of MYC following induction of NFATC4 can partially rescue the quiescent phenotype.

## Results

### NFATC4 mRNA and activity are enriched in a population of slowly dividing CSC

NFAT family members have been linked with quiescence in hair follicle stem cells ^5^. We therefore evaluated the expression of NFAT family members in ovarian cancer stem-like cells (CSC). We previously identified a subset of ovarian CSC marked by expression of ALDH and CD133 ^9^. Evaluation of NFAT family mRNAs in ALDH^+^/CD133^+^ ovarian CSC and ALDH^−^/CD133^−^ ovarian cancer bulk cells identified *NFATC4* as upregulated (4-200 fold, p<0.05-0.001) in 3 independent late stage (III-IV) High-Grade Serous Carcinoma (HGSC) patient-derived ALDH^+^/CD133^+^ samples (Fig. 1a). While not as prominent, *NFATC4* expression is also enriched in slower growing CD133^+^ CSC populations from OVSAHO and A2780 cell lines (cell lines chosen to have distinct CD133^+^ cell populations) (Fig. 1b, c).

**Figure 1.**
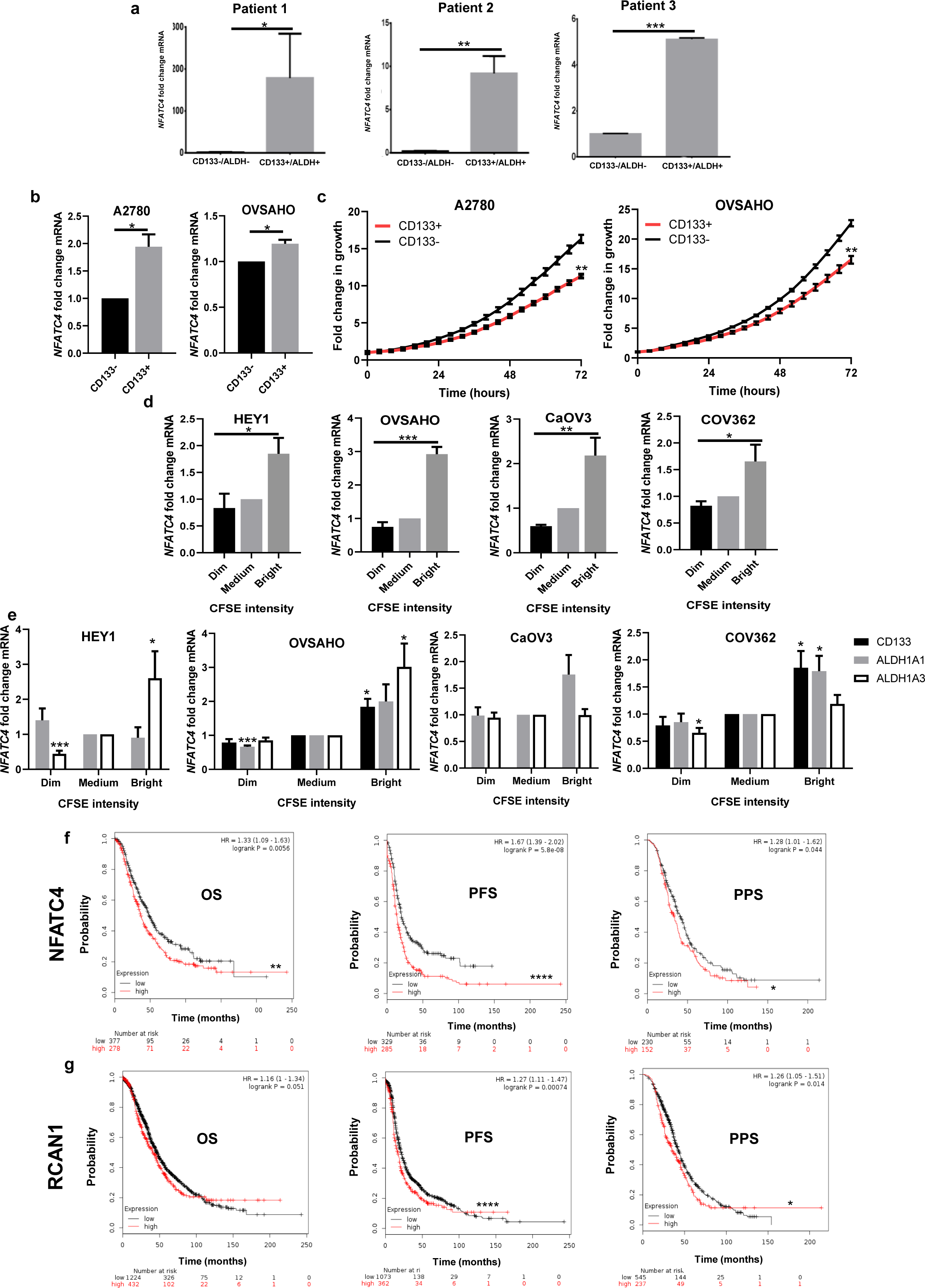
NFATC4 is enriched in ovarian stem like cancer cells and expression correlates with a decrease in cellular proliferation and patient survival. **a** *NFATC4* mRNA expression in ALDH^+^/CD133^+^ ovarian cancer stem-like cells and bulk ALDH^−^/CD133^−^ cancer cells from 3 primary advanced stage (Stage III-IV) HGSC patients. **b** *NFATC4* mRNA expression in CD133^+^ and CD133^−^ ovarian cancer cell lines. **c** Representative growth curves of CD133^+^ and CD133^−^ ovarian cancer cell lines. **d** *NFATC4* mRNA expression levels in 4 cell lines stained with CFSE. CFSE intensity; bright=slowly dividing, medium=bulk cells, dim=rapidly dividing. **e** mRNA expression of the dominate ALDH genes (ALDH1A1/3) and CD133 in CFSE stain cell lines. Kaplan-meier survival plots displaying Overall Survival (OS), Progression Free Survival (PFS), Post Progression Survival (PPS) of TCGA HGSC patients expressing **f** high or low *NFATC4 **g*** high and low *RCAN1*. All experiments were repeated a minimum of 3 times with 3 technical replicates per experiment. T-tests and one-way ANNOVARs were performed to determine significance. *p<0.05, **p<0.01, ****p<0.0001.

To agnostically determine if *NFATC4* was enriched in slower proliferating cells, we used evaluated NFATC4 expression in slowly proliferating/vital dye retaining cells ^13^ in multiple ovarian cancer cell lines. Slow growing/dye retaining cells (Bright), demonstrated a significant enrichment for *NFATC4* mRNA expression compared to the fast-growing (dim – dye dilute) cells in all four cell lines tested (HEY1 p<0.05, OVSAHO p<0.001, CaOV3 p<0.01, COV362 p<0.05) (Fig. 1d). These slow dividing cells were also shown to be enriched for ovarian CSC markers (Fig. 1e).

Suggesting these findings may have clinical relevance, i*n silico* analysis of the impact of NFATC4 expression on patient prognosis was performed using publicly available data ^14,15^. Analyses of microarray data from 1287 HGSC ovarian-cancer patients ^15^ suggested higher expression of *NFATC4* is correlated with worse Overall Survival (OS), Progression Free Survival (PFS) and Post Progression Survival (PPS) (Fig. 1f, p<0.01, p<0.0001, p<0.05). Similarly, analysis of 376 samples in the The Cancer Genome Atlas (TCGA) ovarian cancer data set, demonstrated that dysregulation of the NFATC4 pathway correlated with poor patient outcome (**Supplementary Fig. 1**, p<0.05). Parallel analysis of NFATC4 target gene RCAN1, also showed a correlation between elevated expression and OS, PFS, and PPS (Fig. 1g, p<0.051, p<0.0001, p<0.05). The impact of RCAN on prognosis was less prominent but is likely complicated by RCAN expression in T cells.

### NFATC4 activity induces a quiescent state

To directly interrogate the function of NFATC4 in ovarian cancer cells, we used two distinct previously generated NFATC4 expression constructs, a constitutively active (cNFATC4) ^16^ and an inducible (IcNFATC4) ^17^.NFAT proteins are primarily regulated through phosphorylation regulated cytoplasmic retention (dephosphorylation results in nuclear translocation and activation of transcription various binding partners ^18,19^). One construct (cNFATC4) lacks the regulatory phosphorylation domain and is therefore constitutively nuclear/active (Fig. 2a). Transfection of this construct into A2780 cells demonstrated clear expression of the *NFATC4* mRNA relative to Control-YFP transfected cells (Fig. 2b). Confirming cNFATC4 is transcriptionally active, expression of cNFATC4 resulted in a strong induction of the known NFAT target genes *RCAN1* (regulator of calcineurin-1) ^20^, and Follistatin (FST) ^21^ (Fig. 2b). **To confirm the was broadly applicable, we repeated this experiment and found similar results using** multiple ovarian cancer cell lines (CaOV3, COV362 and OVSAHO) (Fig 2b).

**Figure 2.**
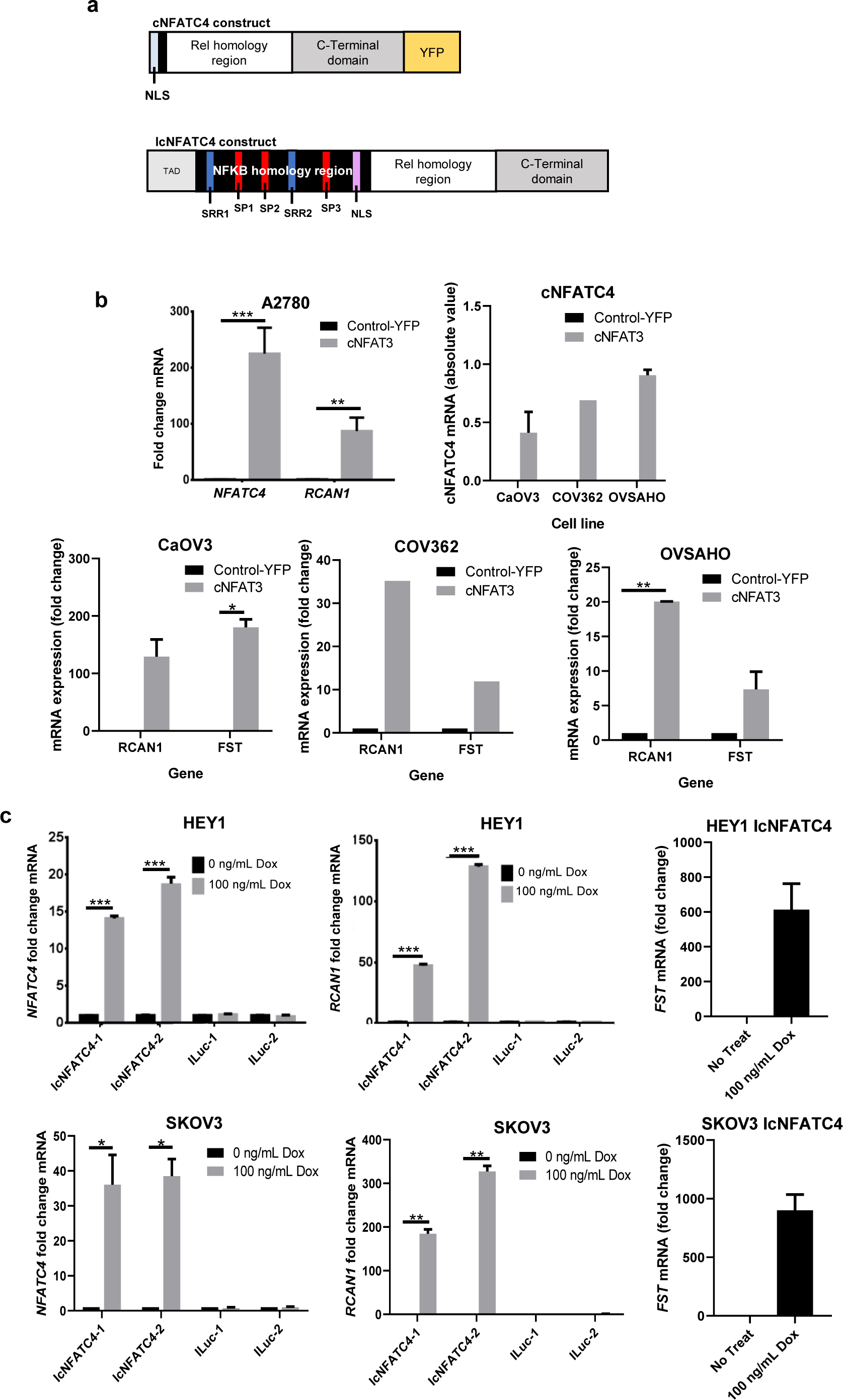
Characterization of NFATC4 overexpression constructs. **a** Schematic diagram of the two constitutive activation *NFTAC4* overexpression constructs; cNFATC4 (truncated regulatory domain and tagged with a yellow fluorescent protein (YFP)), and IcNFATC4 (NLS phosphorylation sites mutated). **b** cNFATC4, RCAN1 and FST mRNA expression in HGSC cells (CaOV3, OVSAHO and COV362) and A2780 cells expressing cNFATC4 or Control-YFP construct (n=2). **d** *NFATC4, RCAN1* and FST (n=3) mRNA expression in HEY1 and SKOV3 cells expressing IcNFATC4 or a ILuc control constructs treated with or without 100 ng/mL doxycycline for 72 h. T-tests and one-way ANNOVARs were performed to determine significance. All experiments were repeated a minimum of 3 times with at least 3 technical replicates per experiment. *p<0.05, **p<0.01, ***p<0.001.

We also generated an inducible nuclear NFATC4 (IcNFATC4) with a puromycin selection cassette. As deletion of the regulatory domain could lead to unexpected changes in function, we used a previously developed construct with point mutations that change the regulatory serines to alanines, leaving the remaining protein intact ^17^. Due to the lack of serine phosphorylation, this NFATC4 protein has an exposed nuclear localization sequence and is therefore constitutively nuclear. An inducible luciferase (ILuc) was used as a control. Disappointingly and inexplicably, despite clear presence of the construct, we were unable to show any inducible expression in multiple HGSC cell lines including OVSAHO, OVCAR3 and OVCAR4 (**Supplemental Fig. 2a**). However, we were able to generate inducible expression of *NFATC4* mRNAs in the HGSC cell line HEY1 ^22^, and the endometriod ovarian cancer SKOV3 line (Fig. 2c). Once again, confirming transcriptional activity, the NFAT target genes *RCAN1* and *FST* were induced following NFATC4 induction (Fig. 2c).

Using these constructs, we tested the effects of NFATC4 activity on ovarian cancer cell growth. Compared to control-YFP lines, cNFATC4 overexpression was associated with a 2-fold (A2780) decrease in total cell number over four days (p<0.0001) (Fig. 3a). Similarly, cNFATC4 overexpression in HGSC cell lines resulted in a 60% (COV362, P<0.001), 50% (OVSAHO, P<0.05) and 70% (CaOV3, P<0.01) decrease in total cell number compared to Control-YFP lines (Fig. 3b).

**Figure 3.**
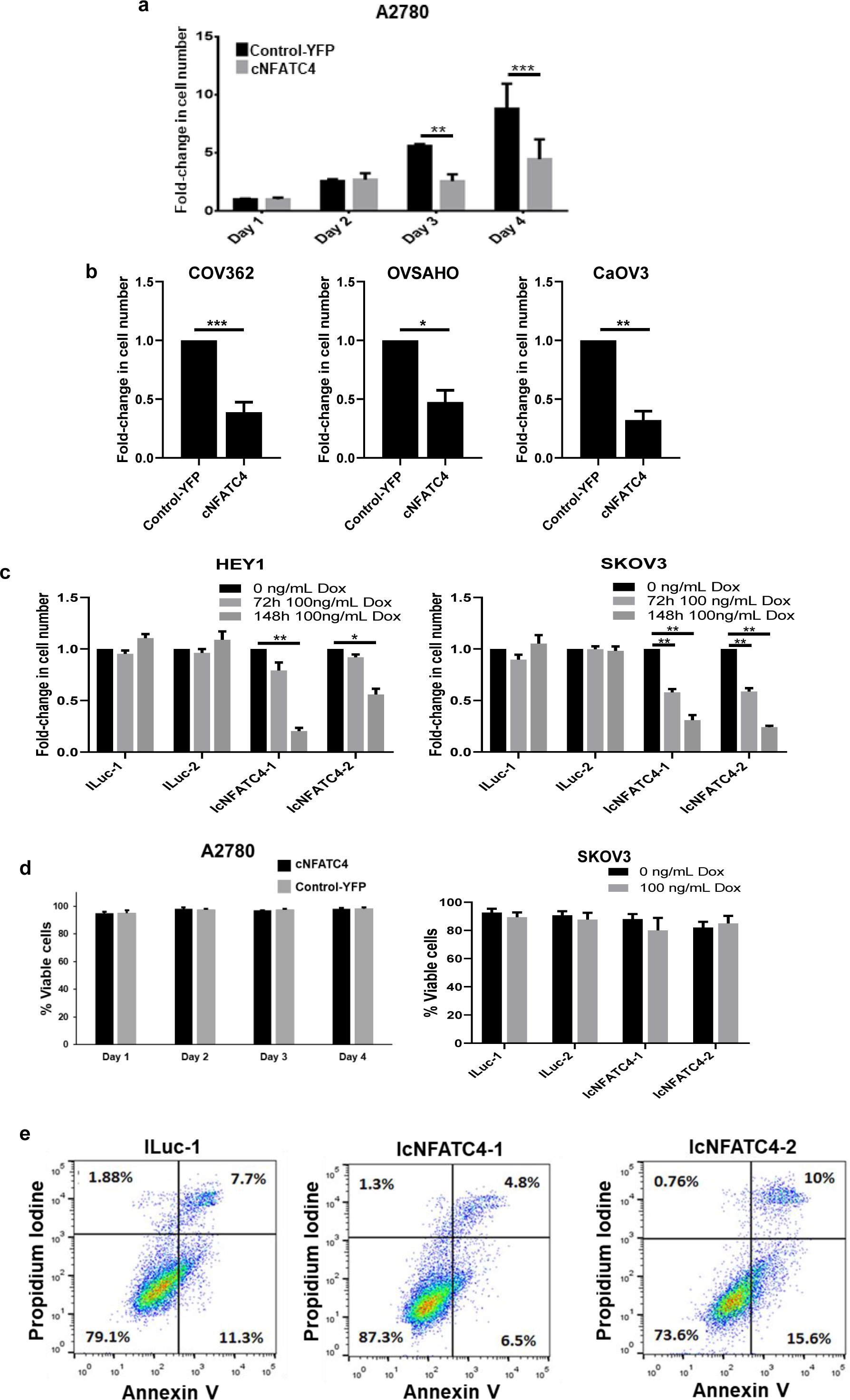
NFATC4 overexpression significantly inhibits cell growth. Cell counts of **a** A2780 cell line or **b** HGSC cell lines (COV362, OVSAHO and CaOV3, at 6, 4 and 6 days respectively) expressing cNFATC4 or control-YFP constructs. **c** Cell counts of HEY1 and SKOV3 cells expressing IcNFATC4 or ILuc control constructs treated with or without doxycycline. **d** trypan blue viability staining of cells line expressing cNFATC4 or YFP control, or IcNFATC4 or ILuc control with or without 100 ng/mL doxycycline. **e** Annexin V/propidium iodine staining of A2780 cells expressing cNFATC4 or YFP control. T-tests and one-way ANNOVARs were performed to determine significance. All experiments were repeated a minimum of 3 times with at least 3 technical replicates per experiment. *p<0.05, **p<0.01, ***p<0.001.

For the IcNFATC4 constructs, doxycycline induction of two independent HEY1 IcNFATC4 cell clones and two independent SKOV3 cell clones showed 1.5-3 fold and 2-4 fold (p<0.01) decrease in cell number at 3 and 6 days after doxycycline treatment, respectively (Fig. 3c). In contrast, doxycycline had no impact on ILuc cell growth for either cell line (Fig. 3c). Explaining the more profound suppression of growth with the IcNFATC4 construct, we noted that, despite multiple rounds of FACS enrichment, cNFATC4 expression was rapidly lost/selected against over time in cell culture which selects for rapidly growing cells (**Supplemental Fig. 2b**).

Given the significant reduction in cell numbers and links between NFAT proteins and apoptosis ^23^, we evaluated the effects of NFATC4 on cellular viability. Trypan blue staining of A2780 and SKOV3 cells indicated that total viability did not change with the expression of cNFATC4 or IcNFATC4 (with or without dox) compared to their respective controls (Fig. 3d). We also analyzed apoptosis rates in the HEY1 IcNFATC4 cells vs ILuc control with Annexin-V FACS. We observed no significant increase in Annexin stain in IcNFATC4 cells vs. ILuc controls (Fig. 3e). Thus, it does not appear that increased apoptosis rates account for the significant reduction in proliferation of NFATC4 overexpressing ovarian cancer cells.

Another explanation for a reduction in growth following overexpression of NFATC4 could be an increase in cellular senescence ^24^. Senescent cells demonstrate an increase in senescence-associated β-galactosidase (SABG). SABG staining demonstrated no increase in SABG expression in control or cNFATC4 cells compared with controls (**Supplementary Fig. 3b**). Therefore, it appears that NFATC4 expression decreases cell division without inducing death, apoptosis, or senescence.

We next evaluated the impact of NFATC4 expression on cellular division. Suggesting a reduction in the percentage of dividing cells, immunofluorescent evaluation of BrdU incorporation demonstrated a reduction of BrdU incorporation in cNFATC4 cells (p=0.058) (Fig. 4a). We then evaluated cellular proliferation at a single cell level. We monitored cell divisions of cells expressing cNFATC4 or Control-YFP in single cell microfluidic culture chips (Fig. 4b), described previously ^11^. Over four days, 39% of control cells and 20% of cNFATC4 cells underwent at least one cell division. 41% of dividing control cells underwent a second cell division while only 4% of the cNFATC4 cells underwent a second division; this resulted in a final >3-fold decreased total cell number in the cNFATC4 cells vs. Control-YFP cells (Fig. 4b). We also evaluated the impact of NFATC4 on growth rates of SKOV3 and HEY1 cell lines. Cells lines expressing the construct IcNFATC4 or ILuc were evaluated in the presence of doxycycline for 96 hours using real-time imaging. Doxycycline treated IcNFATC4 expressing cells had significantly slower proliferation rates than doxycycline treated ILuc controls (HEY1 p<0.0001, SKOV3 p<0.001), with near complete arrest at 96 hours (Fig. 4c) (p<0.0001).

**Figure 4.**
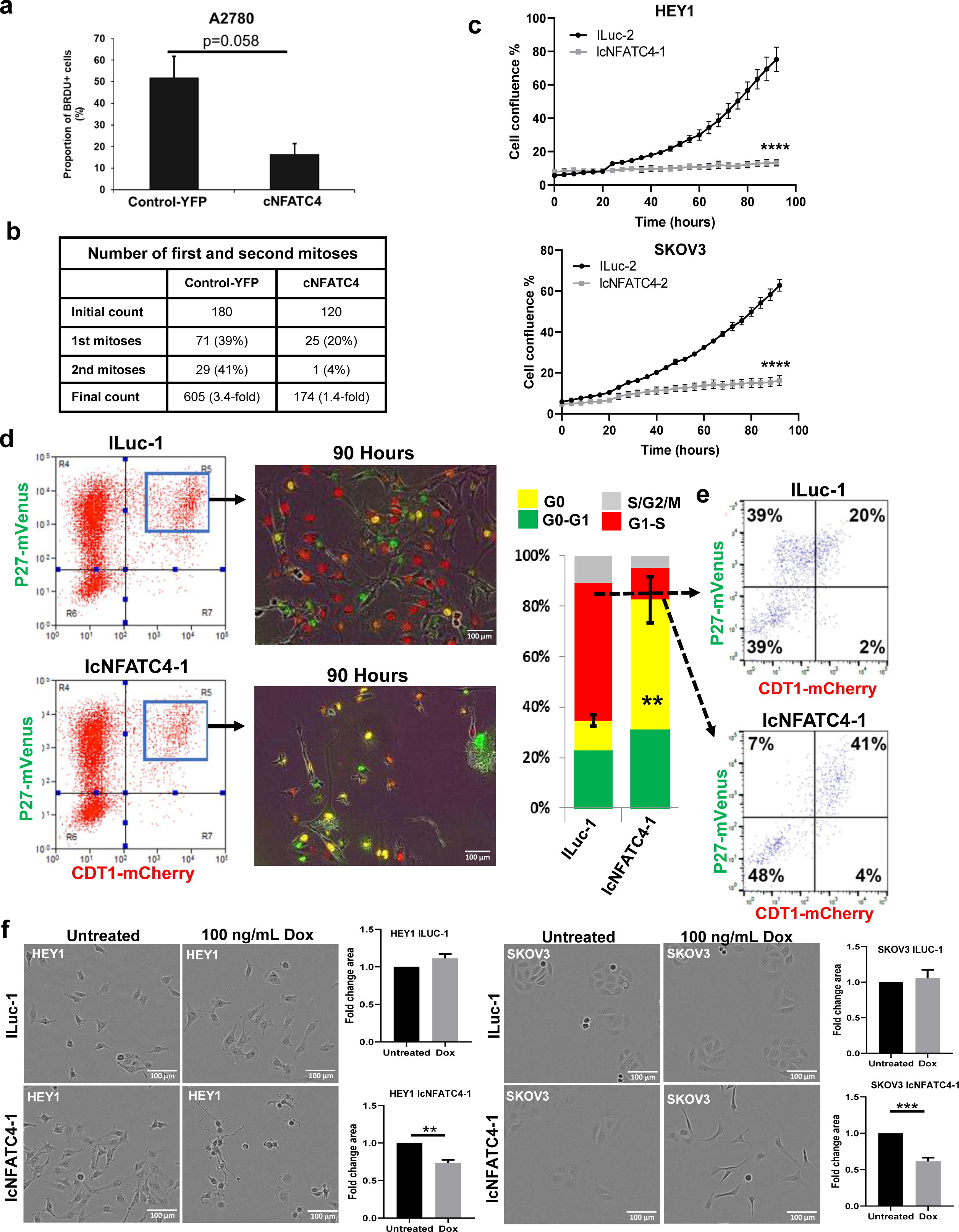
NFATC4 promotes a quiescent phenotype. **a** Quantitation of BrdU incorporation of A2780 cells expressing control-YFP or cNFATC4. **b** Table displaying the number of first and second mitosis of A2780 cells expressing the cNFATC4 or control-YFP constructs, grown in single cell capture microfluidic chips. **c** IncuCyte growth curves of IcNFATC4 or ILuc cells treated with or without doxycycline. **d** FACS plots demonstrating the isolation and quantification of IcNFATC4 and ILuc HEY1 cells expressing the Fucci cell-cycle reporter vectors. Bar graph summarizing the cell cycle phase of cell expressing either construct. **e** Cell cycle analysis of G1-S phase enriched ILuc and IcNFATC4. **f** Representative images and quantification of cells size changes in ILuc and IcNFATC4 cells following doxycycline treatment for 106 h. T-tests and one-way ANNOVARs were performed to determine significance. Microfluidics experiment and Fucci cell cycle experiments were performed twice, all other experiments were repeated a minimum of 3 times with at least 3 technical replicates per experiment. **p<0.01, ***p<0.001.

An explanation for restricted cell proliferation, in the absence of cell death or senescence, is the induction of quiescence. Quiescent cells typically exit the cell cycle and reside in G0. To directly evaluate the impact of NFATC4 activity on G0 cells, we transduced HEY1 cells expressing the IcNFATC4 and ILuc constructs with the mVenus-p27K- and mCherry-CDT1 vectors ^12^. This system can be used to define the G0/G1 transition; briefly, cells expressing high levels of each fusion protein (yellow) are in G0, cells with low or no mVenus-p27K- and high mCherry-CDT1 (red) are transitioning into G1 or in the G1/S phase, and cells with no/low expression of either construct are in S/G2/M. ILuc and IcNFATC4 cells were labeled with the two reporters and the cells expressing both reporter constructs were FACS isolated (Fig. 4d). Purified cells were then treated with doxycycline, plated at ~10% confluency, allowed to adhere overnight before real-time imaging was performed over 90 hours. At the conclusion of the experiment, we scored the number of cells in each phase of the cell cycle found IcNFATC4 cells had a >4-fold increase in the number of cells in G0 (Fig. 4d, P<0.01).

To determine if a subset of cells were cycling while a distinct subset was arresting in G0, we FACS isolated ILuc and IcNFATC4 cells in the G1/S phase of the cell cycle (mCherry-CDT1^(+)^, mVenus-p27K-^(-)^) of the cell cycle, re-plated and FACS analyzed after 72 hours. FACs analysis showed that while ILuc cells redistributed appropriately through the cell cycle with 39% of cells in G1/S, nearly 90% of the IcNFATC4 cells were in the S/G2/M and G0 phases of the cell cycle with only 7% of cells in the G1/S phase of the cell cycle (Fig. 4e).

In addition to G0 arrest, quiescent cells have a unique phenotype which includes a reduction in cell size ^25^. Light microscopic evaluation of doxycycline treated HEY1 and SKOV3 IcNFATC4 and ILuc controls, controlled for cell confluences (**Supplemental Fig 4a**) demonstrated IcNFATC4 cells became significantly smaller with doxycycline treatment (HEY1 p<0.01, SKOV3 p<0.001) (Fig. 4f). FACS analyses of forward-scatter as another measure of size confirmed these results in A2780 cNFATC4 cells and doxycycline treated HEY1 IcNFATC4 cells (**Supplementary Fig. 4b-d**).

### NFATC4 overexpression promotes chemotherapy resistance in vitro

Multiple groups have reported that quiescent/slower-cycling cells are more chemotherapy resistant ^26–28^. We therefore tested the effects of constitutive NFATC4 expression on chemoresistance. We co-treated A2780 cells expressing the cNFATC4 or Control-YFP construct and SKOV3 expressing the IcNFATC4 or ILuc constructs with doxycycline and varying concentrations of cisplatin for 72h (for IC 50 values, see **Supplementary Table 1**). We then quantitated cell number and normalized to the untreated cells. cNFATC4 and IcNFATC4 cells demonstrated significantly increased survival (p<0.001 and p<0.01 respectively) in response to cisplatin chemotherapy when compared to Control-YFP and doxycycline treated ILuc (Fig. 5a). SKOV3 expressing IcNFATC4 treated with and without doxycycline also showed a similar effect (**Supplementary Fig. 5a**, left graph P<0.01), while ILuc untreated vs doxycycline showed no difference (**Supplementary Fig. 5a**, right graph). To confirm these results, we co-treated HEY1 and SKOV3 cells expressing the IcNFATC4 or ILuc control construct with doxycycline and cisplatin (9.5 ug/ml) for 72h and measured cell confluency using the IncuCyte real time imaging (Fig. 5b). IcNFATC4 cells were significantly more resistant to chemotherapy than ILuc cells for both HEY1 (p<0.0001) and SKOV3 (p<0.0001) cell lines.

**Figure 5.**
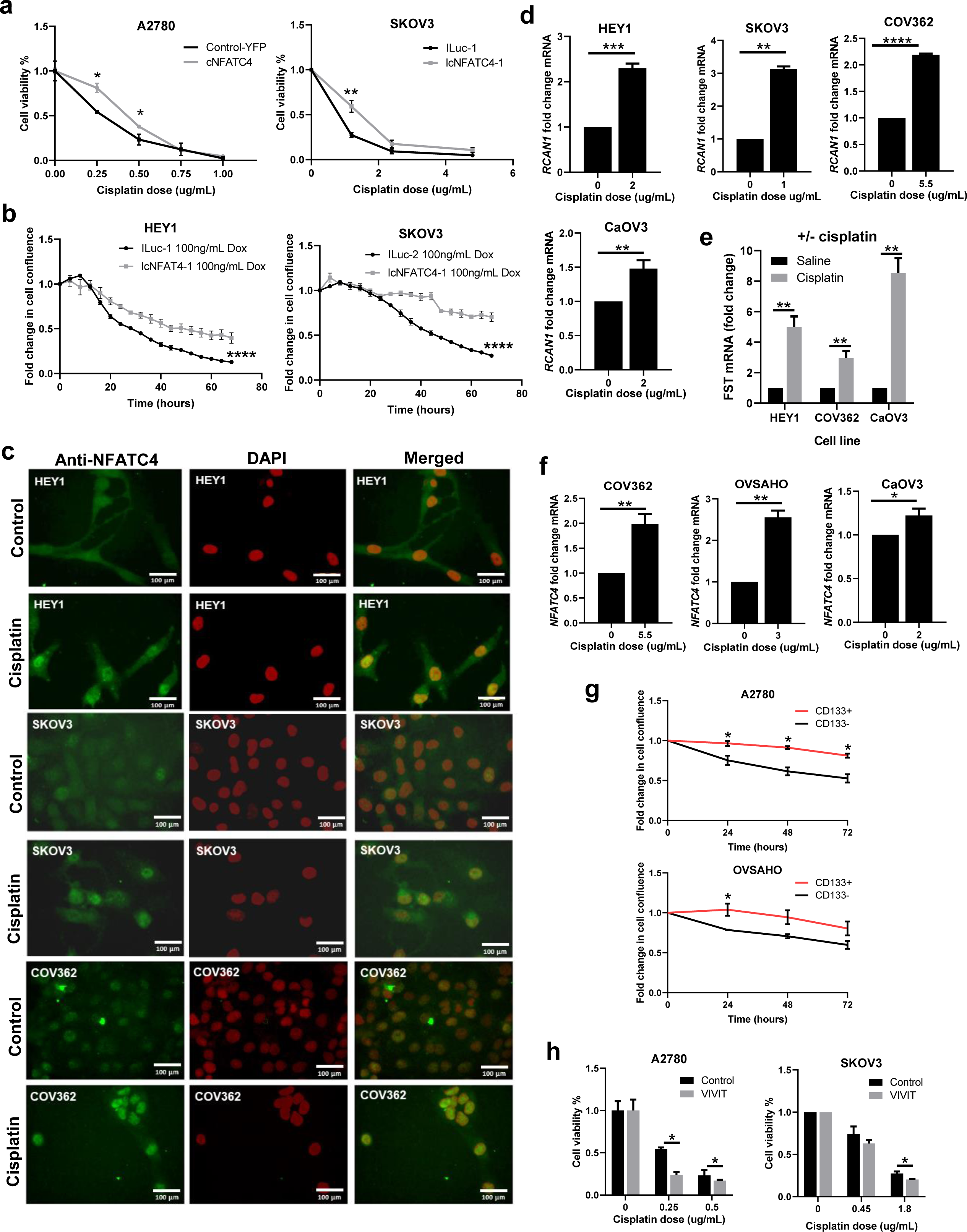
NFATC4 promotes chemoresistance in vitro and is activated by cisplatin. **a** Viability of cells expressing construct pairs (cNFATC4/Control-YFP or IcNFATC4/ILuc) treated with various concentrations of cisplatin. **b** IncuCyte confluence growth curves of IcNFATC4/ILuc expressing cells co-treated with cisplatin and doxycycline. **c** NFATC4 immunofluorescence of cells lines treated with or without cisplatin. **d** *RCAN1* mRNA expression levels of cells treated with or without cisplatin. **e** *FST* mRNA expression levels of treated with or without cisplatin. CaOV3 (2ug/mL), COV362 (5.5ug/mL), HEY (2.5 ug/mL). **f** *NFATC4* mRNA expression of cell lines treated with a high concentration of cisplatin for 72h. **f** IncuCyte confluence growth curves of CD133^−^ vs CD133^+^ cells treated with cisplatin. **g** Cell viability following co-treatment with cisplatin and the pan-NFAT inhibitor VIVIT. T-tests and one-way ANNOVARs were performed to determine significance. All experiments were repeated a minimum of 3 times. *p<0.05, **p<0.01, ***p<0.001.

To determine if chemotherapy exposure could induce NFATC4 nuclear translocation, we performed immunofluorescence on cisplatin treated ovarian cancer cell lines. Cisplatin demonstrated clear nuclear translocation of native NFATC4 in all tested ovarian cancer cell lines (Fig. 5c). Confirming transcriptional activity of NFAT with cisplatin induced nuclear translocation, cisplatin treatment resulted in increases in both *RCAN1* mRNA (COV362 p<0.0001, SKOV3 P<0.01, HEY1 p<0.001, CaOV3 p<0.01) (Fig. 5d), and *FST* mRNA (COV362 p<0.01, HEY1 p<0.1, CaOV3 p<0.01) (Fig. 5e). We also observed a significant enrichment of NFATC4 mRNA expression (COV362 p<0.01, OVSAHO P<0.01, CaOV3 P<0.05), following prolonged treatment of a high dose of cisplatin for 72h (Fig. 5f). Whether this relates to an increase in NFAT gene expression in platinum treated cells, or selection for cells expressing NFATC4 remains to be determined.

Supporting these results, slower growing/NFATC4 enriched CD133^+^ A2780 and OVSAHO CLSCs (Fig. 1c), demonstrated resistance to cisplatin treatment (Fig. 5g). A2780 CD133^+^ cells which had the highest levels of NFATC4 were most cisplatin resistant. To functionally link NFATC4 activity and chemotherapy resistance, we co-treated cell lines with the pan-NFAT inhibitor VIVIT ^29^ and various concentrations of cisplatin for 72 hours. Cells co-treated with VIVIT and cisplatin showed a significant decrease in cell viability when compared to cells treated with cisplatin alone (A2780 p<0.05, SKOV3 p<0.05) (Fig. 5h). Taken together, our *in vitro* data demonstrates NFATC4 promotes quiescence and chemoresistance in ovarian cancer cells and ovarian CSCs in vitro.

With NFATC4 clearly being activated by cisplatin, we wished to investigate if this response was specific to cisplatin or a general response to cellular stress. To test this, we Taxol treated ovarian cancer cell lines for 72h (for IC50 values see, **Supplementary Table 1**) and looked at expression of the same genes (**Supplementary Fig. 6**). We demonstrated a mild increase in both *RCAN1* and *NFATC4* expression following Taxol treatment.

### cNFATC4 expression suppresses tumor growth and drives chemotherapy resistance *in vivo*

We next examined the effect of NFATC4 expression on tumor xenograft growth. A2780 cNFATC4 tumors demonstrated significant growth delay relative to controls (p<0.0001), with essentially no growth for three weeks and with 2/10 cNFATC4 tumors failing to initiate (Fig. 6a). After three weeks, cNFATC4 tumors resumed normal growth; however, suggesting a requirement for loss of cNFATC4 for resumption of growth, analysis of these tumors demonstrated complete loss of cNFATC4 transgene expression (**Supplementary Fig. 7**).

**Figure 6.**
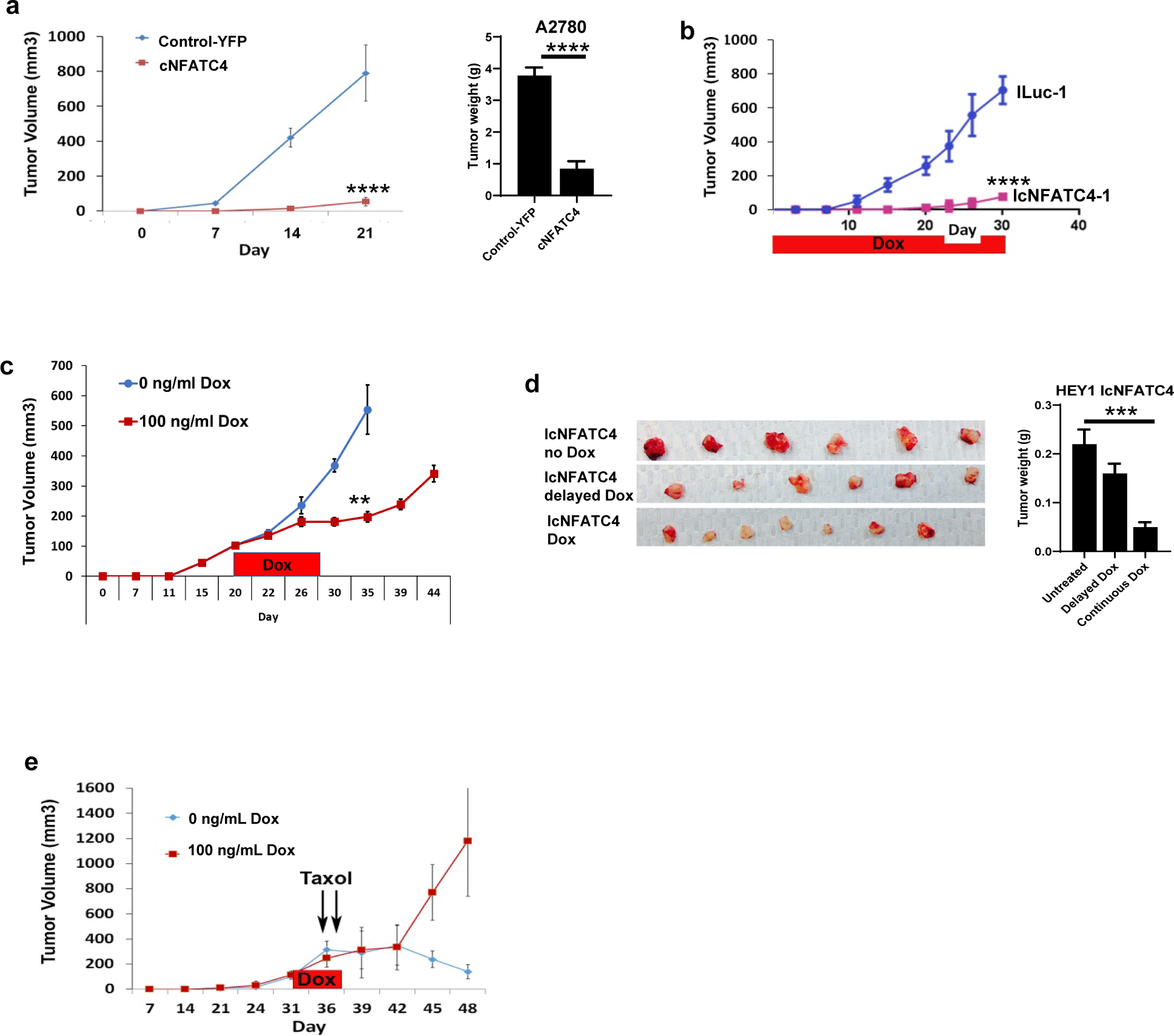
NFATC4 inhibits tumor growth and promotes chemoresistance in vivo. **a** Tumor growth of A2780 cells expressing cNFATC4 or Control-YFP. **b** Tumor growth of HEY1 cells expressing IcNFATC4 or ILuc control constructs in the presences of doxycycline. **c** Tumor growth of IcNFATC4 HEY1 cells treated with delayed doxycycline or vehicle. **d** Tumor weights of HEY1 IcNFATC4 xerographs treated with vehicle, delayed or continuous doxycycline. **e** Tumor growth of HEY1 IcNFATC4 cells treated with doxycycline for 5 days or vehicle, then both treated with 16 mg/kg paclitaxel, intraperitoneally. T-tests and one-way ANNOVARs were performed to determine significance. All experiments were repeated a minimum of 3 times. *p<0.05, **p<0.01, ***p<0.001.

We similarly evaluated the impact of IcNFATC4 expression on tumor growth. In the presence of continuous doxycycline treatment, tumors expressing IcNFATC4 were >13 fold smaller than their ILuc controls (p<0.0001) (Fig. 6b). To confirm this was not related to unequal cell inoculation or altered tumor initiation, we repeated this experiment, but did not initiate doxycycline treatment until tumors were 100mm3. Control-Luc and IcNFATC4 tumors initiated and grew similarly. However, ~5 days after the initiation of treatment with doxycycline, IcNFATC4 tumors showed growth arrest (Fig. 6c-d). Confirming reversibility of the phenotype, on withdrawal of doxycycline, after a slight delay, tumors resumed normal growth.

We next tested the impact of IcNFATC4 induction on chemotherapy resistance. We allowed IcNFATC4 tumors to grow until they were ~150mm^3^. Tumor bearing mice were then randomized and half were then treated with doxycycline for 5 days to induce NFATC4 expression. Due to the low sensitivity of HEY1 to cisplatin at baseline, animals were treated with 2 daily doses of high-dose paclitaxel (16 mg/kg). While control tumors demonstrate a complete response to chemotherapy, tumors in which IcNFATC4 expression was transiently induced demonstrated growth arrest in response to doxycycline, and then ~8 days after doxycycline discontinuation, tumors resumed normal growth without any evidence of response to therapy (p<0.001) (Fig. 6e).

### NFATC4 downregulates MYC and MYC overexpression can partially inhibit early NFATC4 mediated quiescence

It has been reported by multiple studies that NFAT family members can regulate the proto-oncogene MYC ^30–32^. MYC is a master regulator of growth-promoting signal transduction pathways and a well-defined pro-proliferation gene ^33^. To determine if NFATC4, like other family NFAT family members, can regulate MYC expression and if this could be a mechanism for NFATC4 mediated quiescence, we examined the effect of cNFATC4 on MYC mRNA expression. HEY1 and SKOV3 cells demonstrated a significant reduction in MYC expression following NFATC4 induction, p<0.0001 and p<0.05 respectively (Fig. 7a). Furthermore, we conducted doxycycline recovery experiments, where we induced the construct by treating cells with doxycycline for 72h, then removed the doxycycline and recorded cell number (**Supplementary Fig 8**) and mRNA expression of NFATC4 target genes (**Supplementary Fig 9**) as the cells resumed cell cycle. We were then able to re-induce the quiescent state via additional doxycycline treatment. These experiments demonstrated MYC, along with the NFATC4 target genes RCAN1 and FST, cycled with the induction and loss of quiescence, supporting their role in inducing this quiescent state.

**Figure 7.**
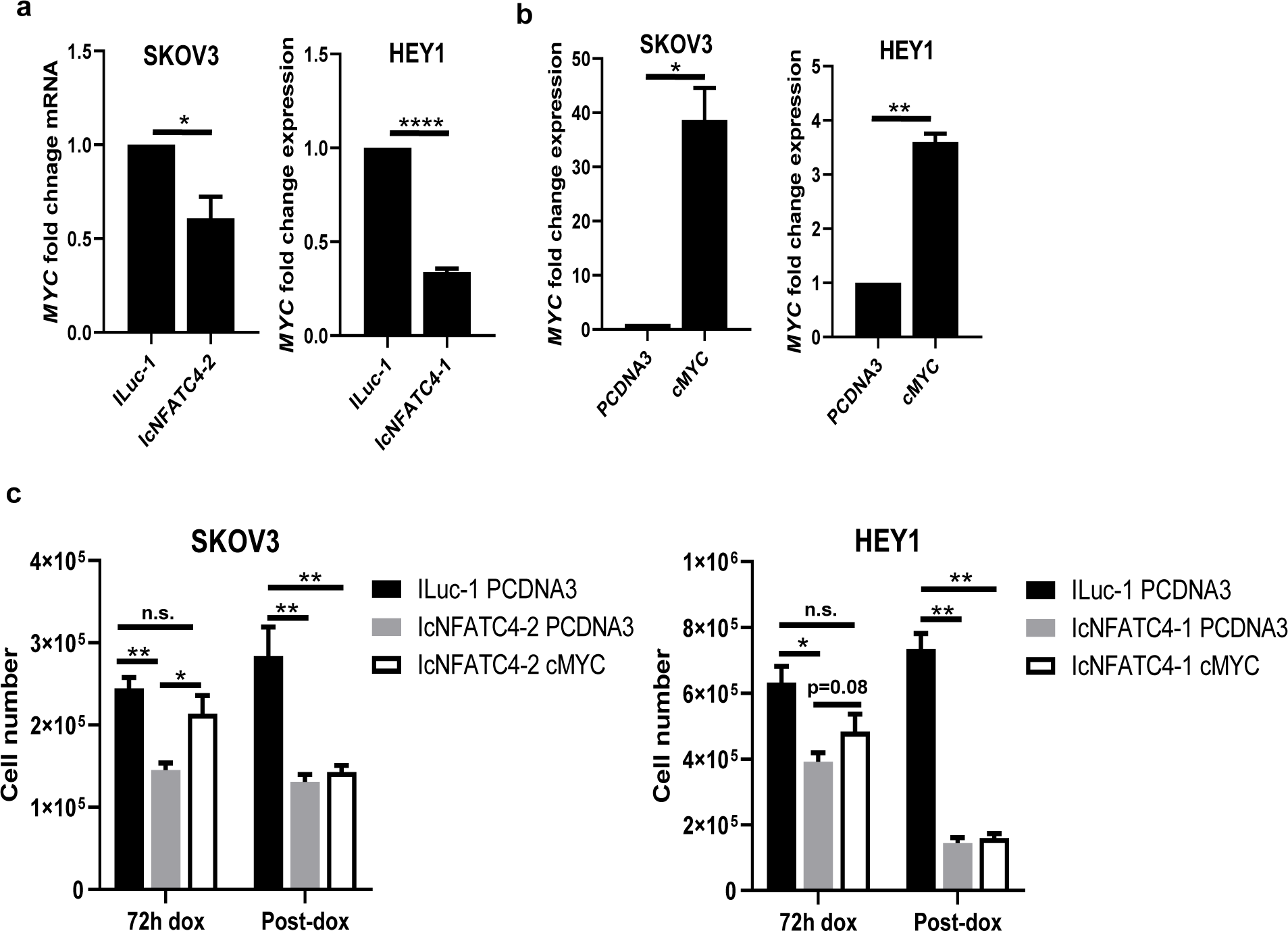
NFATC4 overexpression inhibits MYC, and cMYC overexpression partially rescues the quiescent phenotype at early, but not late, timepoints. **a** MYC mRNA expression following 24 h doxycycline treatment. **b** Validation of cMYC overexpression construct in SKOV3 and HEY1 cell lines. **c** The effect of MYC overexpression on cell number of IcNFATC4 cells transfected with PCDNA3 or cMYC and treated with doxycycline for 72 h or post treated for 72h with dox before transfection with PCDNA3 or cMYC and treated with doxycycline for an addition 72 h. T-tests and one-way ANNOVARs were performed to determine significance. All experiments were repeated a minimum of 3 times. n.s.= not significant, *p<0.05, **p<0.01, ****p<0.0001.

To investigate if this downregulation in MYC following NFATC4 induction could affect cell proliferation, we transfected lcNFATC4 expressing SKOV3 and HEY1 cells with a MYC overexpression construct ^34^ or PCDNA3 vector control. MYC overexpression resulted in a significant increase in MYC mRNA expression (SKOV3 p<0.05, HEY1 p<0.01) (Fig.7b). To determine if early induction of MYC expression was able to prevent NFATC4 induced quiescence, induced IcNFATC4 cells for 6 hours then transfected cells with PCDNA3 (control) or cMYC. We continued doxycycline for 72 h before and then evaluated cell growth. Cells transfected with MYC 6 h after NFATC4 induction demonstrated cell growth which was not statistically significantly different from control iLuc cells (Fig.7c).

To determine if MYC expression could overcome growth suppression in established NFATC4 driven quiescent cells we repeated the above experiment, in cells in which NFATC4 had been induced for 72 h to ensure the cells had already establish a quiescent phenotype before transfecting with PCDNA3 or cMYC. Doxycycline treatment was continued for an additional 72 h before and cell proliferation evaluated. Interestingly, MYC expression in established quiescent cells was unable reverse the quiescent phenotype (Fig.7c).

## Discussion

The Nuclear Factor of Activated T-Cells (NFAT) family of transcription factors act as master regulators of numerous cellular processes. In normal cells, NFAT family members can impact both proliferation ^35–39^ and quiescence ^5^. NFAT proteins have been directly implicated in the regulation of stem cell proliferation ^40^. We report here a critical role for NFATC4 in the regulation of cellular quiescence in ovarian cancer. Expression of constitutively nuclear NFATC4 suppresses cellular proliferation, reduces cell size and arrested tumor growth. Furthermore, consistent with previous reports ^41,42^, we find that NFATC4 activity contributes to chemotherapy resistance, both *in vitro* and *in vivo*. Finally, NFATC4 induction represses MYC which contributes to a quiescent state.

### NFAT family members and regulation of quiescence

We identified NFATC4 as a potential regulator of quiescence in ovarian cancer. Previously NFATC1 was identified as regulating quiescence in the hair follicle. More recently, a study of NFATC3 in the brain ^43^ observed significant changes in cell proliferation and vital dye retention with NFATC3 inhibition. These studies implicate the NFAT family of transcription factors as regulators of quiescence.

The complete mechanism through which NFAT regulates proliferation/quiescence is unclear. In the hair follicle, NFAT regulates the cyclin dependent kinase CDK4 to arrest cell cycle progression ^5^. We did not observe similar mechanisms here; however, we did observe NFATC4 expression correlated with a down regulation of MYC, while MYC overexpression was able to partially rescue the quiescent phenotype following early, but not late, induction of NFATC4. This observation suggests that although downregulation MYC could be an important part of the induction of a quiescent phenotype, there are secondary mechanisms downstream of NFATC4 which are critical in the maintenance of the quiescent state. Future work is required to elucidate the mechanism responsible for NFATC4 induced quiescence.

### NFATC4 and chemotherapy resistance

NFATC4 has been poorly studied in cancer. In normal physiologic states, NFATC4 appears to function partly as a general stress response protein, as it serves a protective role in cardiomyocytes in response to radiation ^44^, is activated by mechanical stress in the heart ^45^ and bladder ^46^, and serves as a protective factor during hypoxia ^47^. NFATC4 may serve as a similar stress regulator in cancer cells to promote survival. We have shown that NFATC4 translocates to the nucleus and initiates transcription in response to cisplatin chemotherapy and drives chemotherapy resistance. Similarly, NFATC4 expression was linked with therapeutic response in gastric cancer ^48^. Taxol’s partial induction the NFATC4 pathway of may be a result of its limited ability to increase intracellular calcium ^49^, while increasing intracellular calcium is a key mechanism by which cisplatin functions ^50^.

Consistent with a role for NFATC4 in cancer stem-like cells, NFATC4 plays a role in pancreatic cell plasticity and tumor initiation ^51^. Consistent with the data above, survival data demonstrating dysregulation or high expression of NFATC4 leads to worse prognosis of ovarian cancer patients, suggest NFATC4 may be an important therapeutic target in ovarian cancer to overcome the chemotherapy resistance associated with slow-cycling cells. This hypothesis is supported by studies on cyclosporine, a commonly used immunosuppressant which inhibits the NFAT family. Cyclosporine has shown activity as a chemo-sensitizer in a phase II clinical trial, demonstrating that cyclosporine could improve response to therapy in patients with chemotherapy-refractory disease ^52,53^. However, this has not been reproducible ^54^ and has not been tested in patients with chemotherapy naïve disease, who may benefit the most from the elimination of slow cycling cells. Furthermore, as cyclosporine impacts all NFAT family members, and is not selective for cancer cells, suppression of immunity may limit efficacy. NFATC4 is the only core NFAT family member that is not expressed in the immune system ^55^, thus the development of specific NFATC4 inhibitors could allow chemosensitization of the NFATC4-expressing CSC without concomitant immunosuppression.

In summary, we have found that the master transcriptional regulator NFATC4 induces a quiescent state in ovarian cancer and translocates to the nucleus in response to chemotherapy. Constitutively nuclear NFATC4 is associated with a reduction in cellular size and proliferation and the induction of chemotherapy resistance. NFATC4 promotes the quiescent phenotype by early downregulation of MYC; however other mechanisms are responsible for maintaining an established quiescent phenotype. Taken together, this data suggest NFATC4 is an important therapeutic target in ovarian cancer that warrants significant further investigation.

## Materials and Methods

### Cell Culture

The A2780 cell line was obtained from Dr. Susan Murphy at Duke University. SKOV3, CaOV3 and HEY1 lines were purchased from ATCC (2018). OVSAHO cells were gifted from Dr. Deborah Marsh from the University of Sydney. COV362 were purchased from Sigma-Aldrich (2018). All cells were cultured in RPMI-1640 media with 10% FBS, 1% Penicillin/Streptomycin at 37°C and 5% CO_2_.

### Constructs

Both constructs used in this study were designed to result in constitutively nuclear NFATC4. A constitutively nuclear NFATC4-YFP fusion (cNFATC4) with the phospho-regulatory domain deleted or a YFP-only control (Control-YFP) were cloned into a pGIPZ lentiviral vector and transduced into the A2780, CaOV3, OVSAHO and COV362 ovarian cancer cell lines. A second, phospho-specific mutant constitutively active NFATC4 ^17^ was also cloned into the doxycycline-inducible Tet-One expression system (Clontech) to create an inducible and constitutive NFATC4 (IcNFATC4) in the HEY1 ovarian cancer line. This was paired with an inducible luciferase control (ILuc) to control for overexpression. Details regarding the structure and validation of all constructs are presented in the results section (Fig. 2). Because the only known function of NFAT proteins are transcription, the phospho-mutants used are constitutively nuclear and therefore constitutively active. A pcDNA3-cmyc construct was purchased from Addgene (Cat# 16011).

### Patient samples

Fresh High-grade Serious Ovarian Carcinomas (HGSOC) were acquired from the University of Michigan’s Comprehensive Cancer Center. Fresh tumor samples were dissociated using a Tumor Dissociation Kit (Miltenyi Biotec) and cultured using standard conditions. HGSOC diagnosis was confirmed using immunohistochemistry.

### Carboxyfluorescein succinimidyl ester (CFSE) Assay

HEY1, COV362, OVSAHO and CaOV3 cell lines were stained with 2.5 uM of CellTrace™ CFSE Cell Proliferation Kit (Thermo Fisher) for 20 mins at 37°C, washed and then grown for 5-7 days. The top 4% bright cells, 10% medium cells and bottom 4% dim cells were FACS isolated. RNA was extracted, and qPCR performed to validate NFATC4 expression (as below).

### Quantitative PCR

RNA was extracted using RNeasy Mini Kit (Qiagen) and cDNA was made using SuperScript™ III Reverse Transcriptase kit (Theremofisher). qPCR was performed using SYBR™ Green PCR Master Mix (Applied Biosystems) using standard cycling conditions. The primers used for this study are available in supplemental material (**Supplementary Table 2**).

### Cell counting

Cell counts were performed using the Moxi Z automated counting system (ORLFO Technologies) and the Cassettes Type S. For trypan blue staining, the Countess automated cell counter (Invitrogen) was used with Countess Cell Counting Chamber Slides.

### BrdU labeling

A2780 cells were treated with 10 µM BrdU labeling solution and incubated for 4 hours at 37ºC in a CO2 incubator. Cell were washed with PBS, fixed with 4% paraformaldehyde and permeabilized with 0.1% Triton X. Standard ICC was followed. Cells were then incubated in 2.5 M HCL for 30 mins at room temperature, then washed with PBS. Standard immunocytochemistry protocol was then followed.

### Micro Fluidics

A2780 Control-YFP or cNFATC4 cells were loaded into single cell capture microfluidic chips from collaborator Dr. Yoon (University of Michigan). Cells mitoses were tracked using a microscope over a three-day period and results were recorded. Cell viability was confirmed using LIVE/DEAD cell staining at the termination of the experiment.

### Size Analysis

HEY1 and SKOV3 cells expressing the ILuc and IcNFATC4 constructs were pre-treated with doxycycline, then plated into a 96 well plate at 300 and 1000 cell/well respectively and placed into the IncuCyte. Cells were treated with or without doxycycline and grown for 36 h. Images we taken at 36 hours making sure the confluency was comparable between the doxycycline treated and untreated control cells (**Supplementary Fig. 4a**). Images were imported into ImageJ and cell size was calculated by drawing around cells and quantifying their area.

### Fluorescence ubiquitination cell cycle indicator (FUCCI)

HEY1 cells expressing the IcNFATC4 or ILuc constructs were transduced with the p27-mVenus and CDT1-mCherry FUCCI cell cycle reporter constructs ^12^. Cells expressing both constructs were isolated using FACS and plated in the IncuCyte and grown for 90 hours. Fluorescence was measured, and percentages of green, red, yellow, and unstained cells were quantitated. G1-S phase cells expressing the constructs were FACS isolated and grown for 3 additional days to determine if they retained the cell cycle phases as a result of the NFACTC4 overexpression.

### Annexin V staining

For apoptosis detection via annexin staining, HEY1 ILuc and IcNFATC4 cells were grown in 6 well dishes with or without doxycycline for 72 hrs. Cells were stained with the Annexin-V FITC apoptosis kit (BD Biosciences) according to the manufacturer’s instructions and at least 10,000 events were analyzed on the Mo Flo Astrios flow cytometer (BD Biosciences). The percentage of Annexin V^+^, PI^+^, Annexin V^+^/PI^+^, and Annexin V^−^/PI^−^ cells was quantified.

### Senescence-associated beta galactosidase (SABG) staining

Cells were plated on tissue culture coverslips, allowed to grow for 96 hours, fixed, and then SABG staining was done as previously described ^56^.

### IncuCyte growth curves

HEY1 and SKOV3 cells expressing the ILuc or IcNFATC4 constructs were seeded at 300 and 1000 cells per well respectively, in a 96 well plate. For growth curves, cells were treated with or without 100 ng/mL doxycycline for 96 h. For cisplatin curves, cells were treated with 9.5 ug/mL cisplatin for 72h with or without co-treatment with 100 ng/mL dox. IncuCyte images were taken every 4 h and cell confluence was recorded.

### Immunofluorescence

HEY1, COV362 and SKOV3 ovarian cancer cell lines were grown on glass coverslips and treated with various concentrations of cisplatin based on their respective IC50s. Coverslips were fixed with 4% paraformaldehyde and permeabilized with 0.1% Triton X. Cells were blocked with 10% horse serum and incubated with 1:50 mouse anti-NFATC4 antibody (Novus Biologicals) in 5% horse serum for 2 hours. Slides were washed 3 x 5 mins with PBS, cells were incubated with an Alexa Fluor 488 anti-mouse secondary antibody, mounted with DAPI mounting medium (Vector Labs), and then imaged on an Olympus BX41 microscope.

### In vivo xenografts

NOD/SCID/IL2R^KO^ or nude mice were injected with 500,000 Control-YFP or cNFATC4 cells or 300,000 ILuc or IcNFATC4 cells for tumor xenograft experiments. Animals were maintained at 12 hour light/dark cycles under SPF conditions with free access to food and water. For induction, 2 mg/mL doxycycline was administered in the water along with 5% sucrose to mask its bitter taste. Tumors were monitored once a week initially and twice a week after tumors reached 1000 mm^3^, and animals were sacrificed at protocol endpoints. All experiments were conducted in accordance with the animal care and use committee from the University of Michigan.

### Statistical analysis

Statistically analysis was conducted using GraphPad Prism (8.0.2) and http://vassarstats.net/. All data was analyzed using two tailed student’s t-tests or one-way ANNOVARs. A minimum of 3 replicate experiments (n≥3) were used for each experiment.

## Acknowledgements

Funding for this work was provided by Ann and Sol Schreiber Mentored Investigator Award (599997) from the Ovarian Cancer Research Alliance (OCRA), DOD Award #W81XWH-15-1-0083 and NIH RO1 award 1R01CA203810.

## Contributions

A.C and M.I conducted the majority of the experiments. R.B and S.B conceived the project and planned the experiments. P.H, E.Y and D.H. helped with the experiments. G.D., K.A., E.Y reviewed the manuscript. A.C, M.I and R.B. wrote the manuscript.

## Supplementary Figure legends

**Supplementary Figure 1.** Kaplan-Meier survival plot. Survival probability of dysregulation of the NFATC4 pathway in ovarian cancer TCGA data set. *p<0.05.

**Supplementary Figure 2.** Evaluation of NFATC4 constructs. **a** NFATC4 and RCAN1 mRNA expression of 3 HGSOC cells lines transduced with IcNFATC4 or ILuc control. **b** Flow cytometry analysis of A2780 cells expressing Control-YFP or cNFATC4 constructs.

**Supplementary Figure 3.** No difference in apoptosis or SABG staining between NFATC4 and control constructs. **a** Annexin V PI apoptosis assay of HEY1 expressing ILuc or NFATC4 constructs. **b** SABG staining of A2780 cNFATC4 or Control-YFP. **c** Quantification of SABG staining; the cyclin D1/CDK4 and CDK6 inhibitor Ribociclib was used as a positive control for senescence.

**Supplementary Figure 4.** lcNFATC4 expressing cells have a reduction in cell size. **a** IncuCyte growth curves of IcNFATC4 or ILuc HEY1 and SKOV3 cells treated with or without doxycycline. Red box indicates the confluency used to validate changes in cell morphology. Forward scatter flow cytometry of; **b** A2780 cNFATC4 vs Control-YFP cells, **c** HEY1 ILuc and lcNFATC4 cells treated with or without doxycycline. **d** Quantitation HEY1 cell size. **p<0.01. ****p<0.0001.

**Supplementary Figure 5.** NFATC4 promotes chemoresistance. Cell viability ILuc and IcNFATC4 SKOV3 cells treated with or without doxycycline and co-treated with varying concentrations of cisplatin.**p<0.01.

**Supplementary Figure 6.** Taxol partially activates NFATC4 signaling in ovarian cancer cell lines. OVSAHO, COV362 and HEY1 cells were treated with or without 3 uM and 9 uM of Taxol for 72 h before RCAN1 and NFATC4 expression was validated using qPCR. Results were expressed as fold change to untreated control, n=2.

**Supplementary Figure 7.** NFATC4 constructs expression is lost in vivo. Flow cytometry analysis of control-YFP and cNFATC4 expression in tumors.

**Supplementary Figure 8.** Doxycycline recovery cell counts of NFATC4 expressing SKOV3 and HEY1 cells lines. **a** HEY1 and SKOV3 cells expressing the IcNFATC4 and ILuc constructs were treated for 72h with 100 ng/mL doxycycline. After 72h doxycycline was removed and cells were grown in doxycycline free media for 120-144 h. Cell counts were performed daily. **b** after recovering from quiescence, cells were treated with dox for 96 h, then re-plated at 30 K cells and grown in the presence of dox for an additional 72-96h. Cell counts were recorded daily. n=2.

**Supplementary Figure 9.** Doxycycline recovery qPCR of NFATC4 target genes in NFATC4 expressing SKOV3 and HEY1 cells lines. HEY1 and SKOV3 cells expressing the IcNFATC4 and ILuc constructs were treated for 72h with 100 ng/mL doxycycline. After 72h doxycycline was removed and cells were grown in doxycycline free media for 120 h. After recovering from quiescence, cells were treated with dox for another 144h. RNA was extract and qPCR performed for NFATC4, FST, RCAN1 and MYC at each of the listed time points. N=2.

**Supplementary Table 1.** Cisplatin and Taxol IC50 values for ovarian cancer cell lines used in this study.

**Supplementary Table 2.** qPCR primer sequences.

