## Supplementary data for "NFATC4 Promotes Quiescence and Chemotherapy Resistance in Ovarian Cancer" (from another version of this preprint: version 1, 2019-10-31)

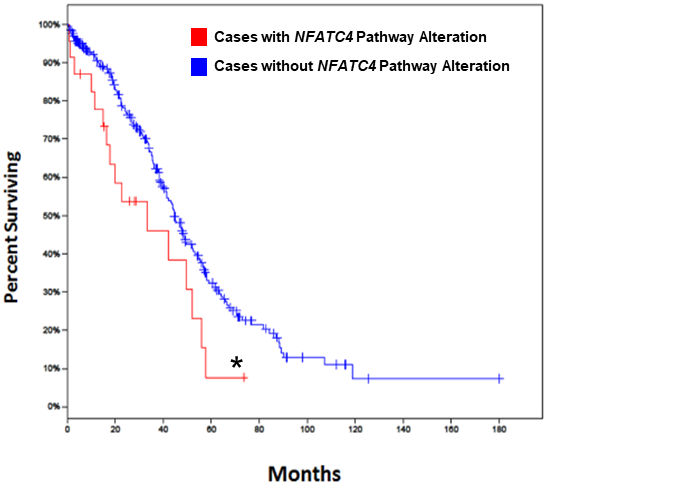

**Supplementary Figure 1**

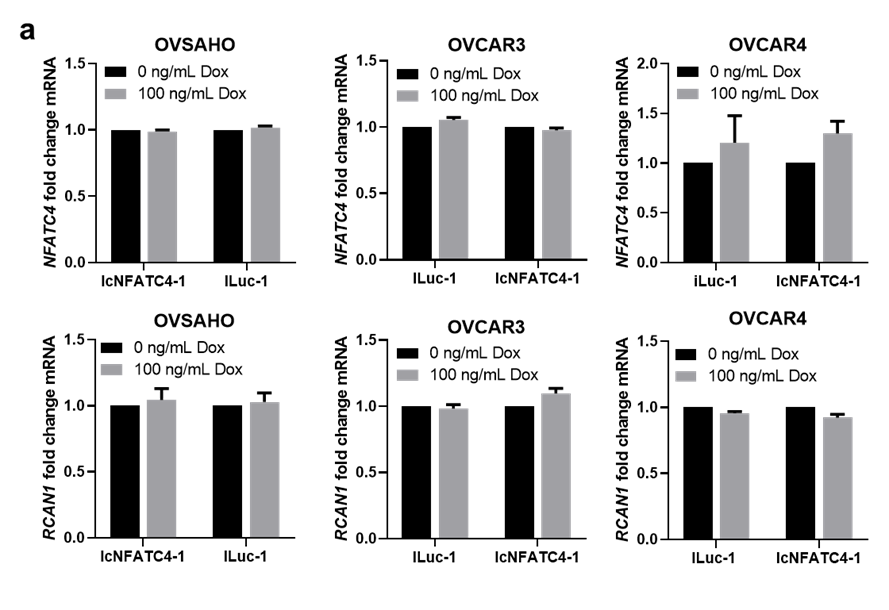

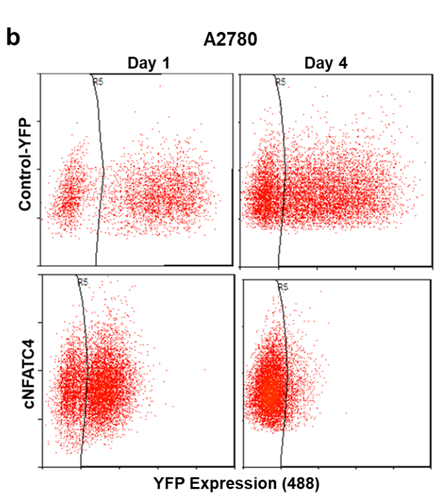

**Supplementary Figure 2**

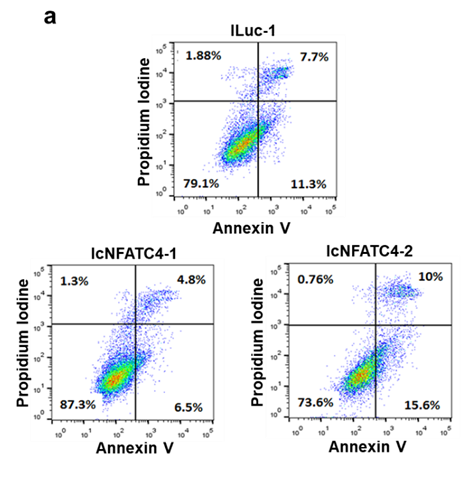

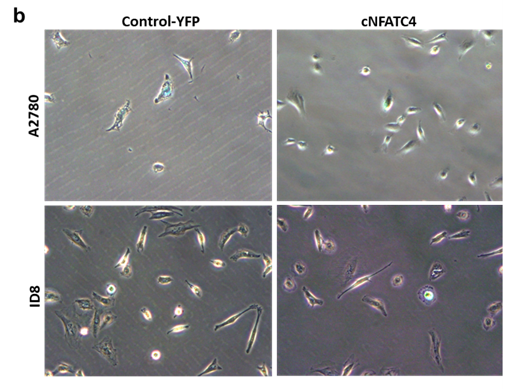

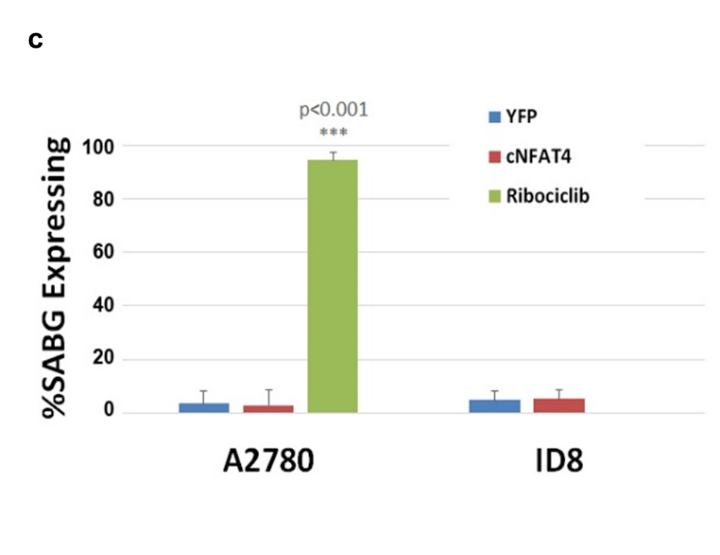

**Supplementary Figure 3**

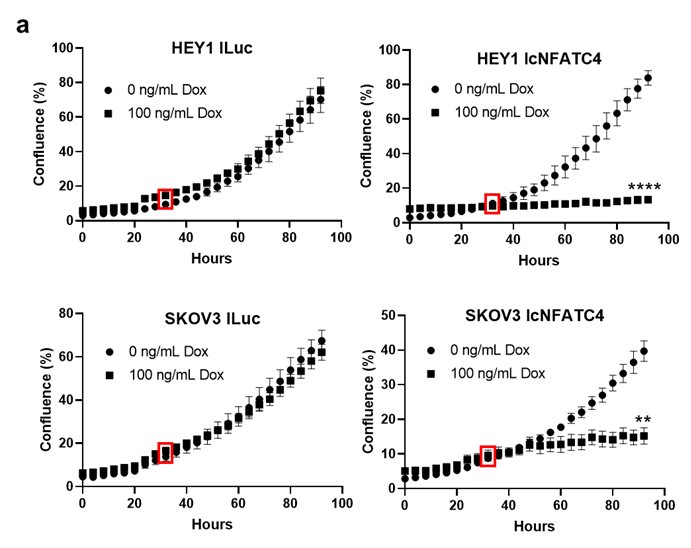

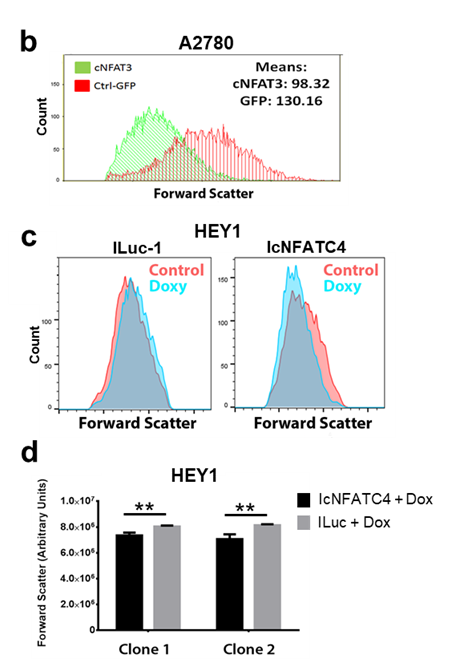

**Supplementary Figure 4**

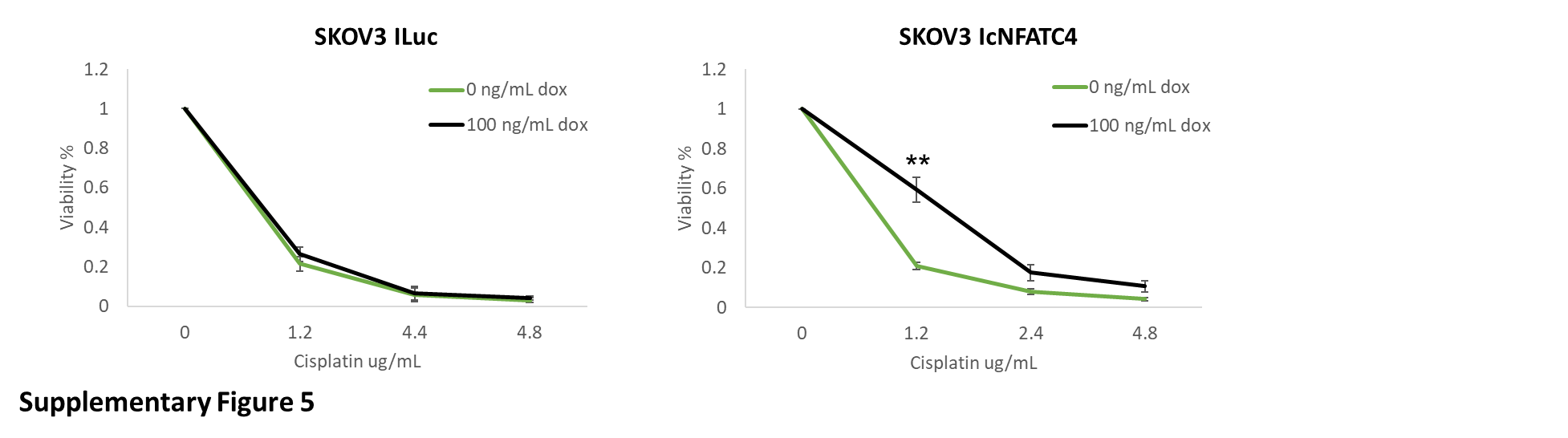

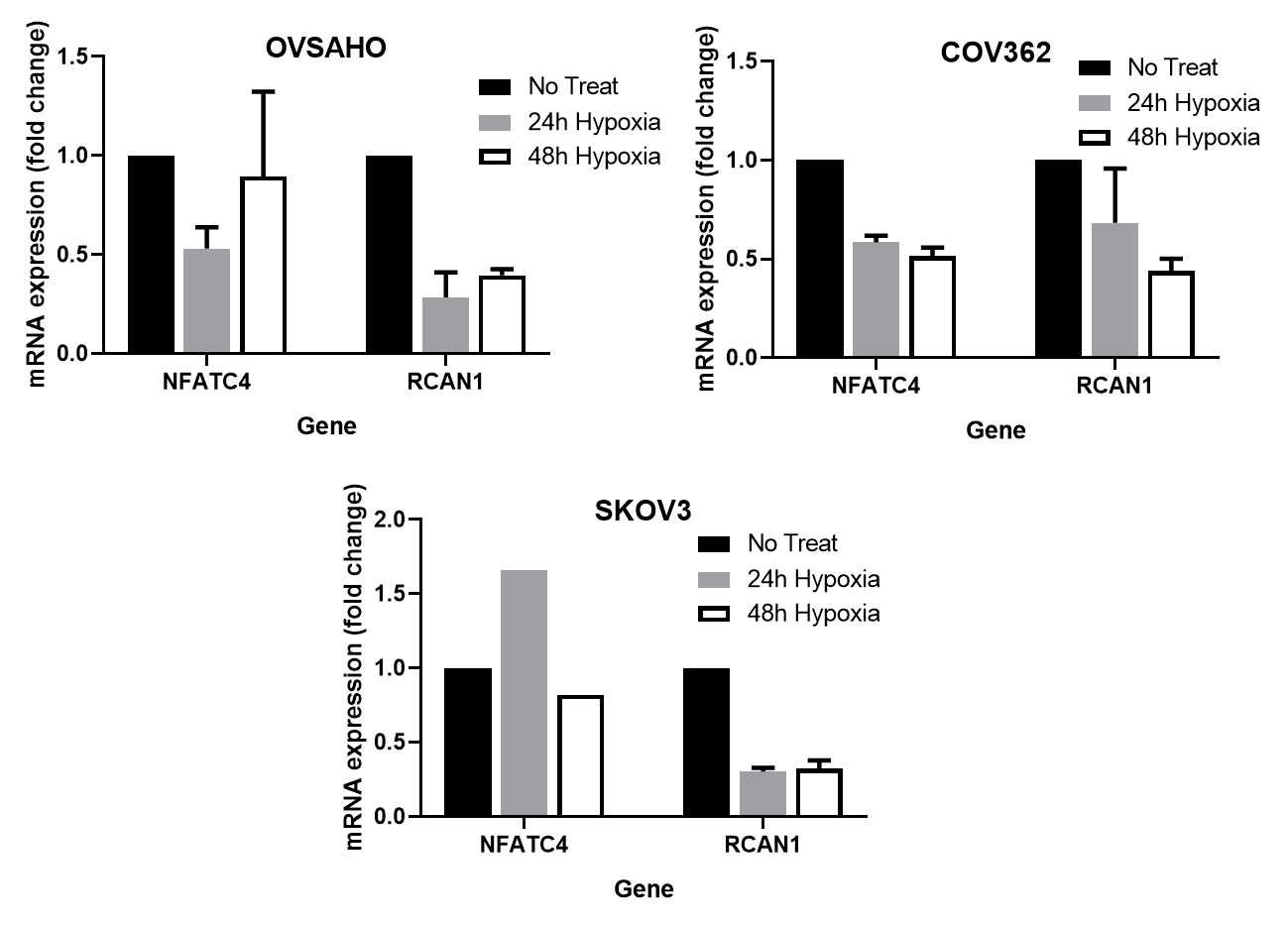

**Supplementary Figure 6**

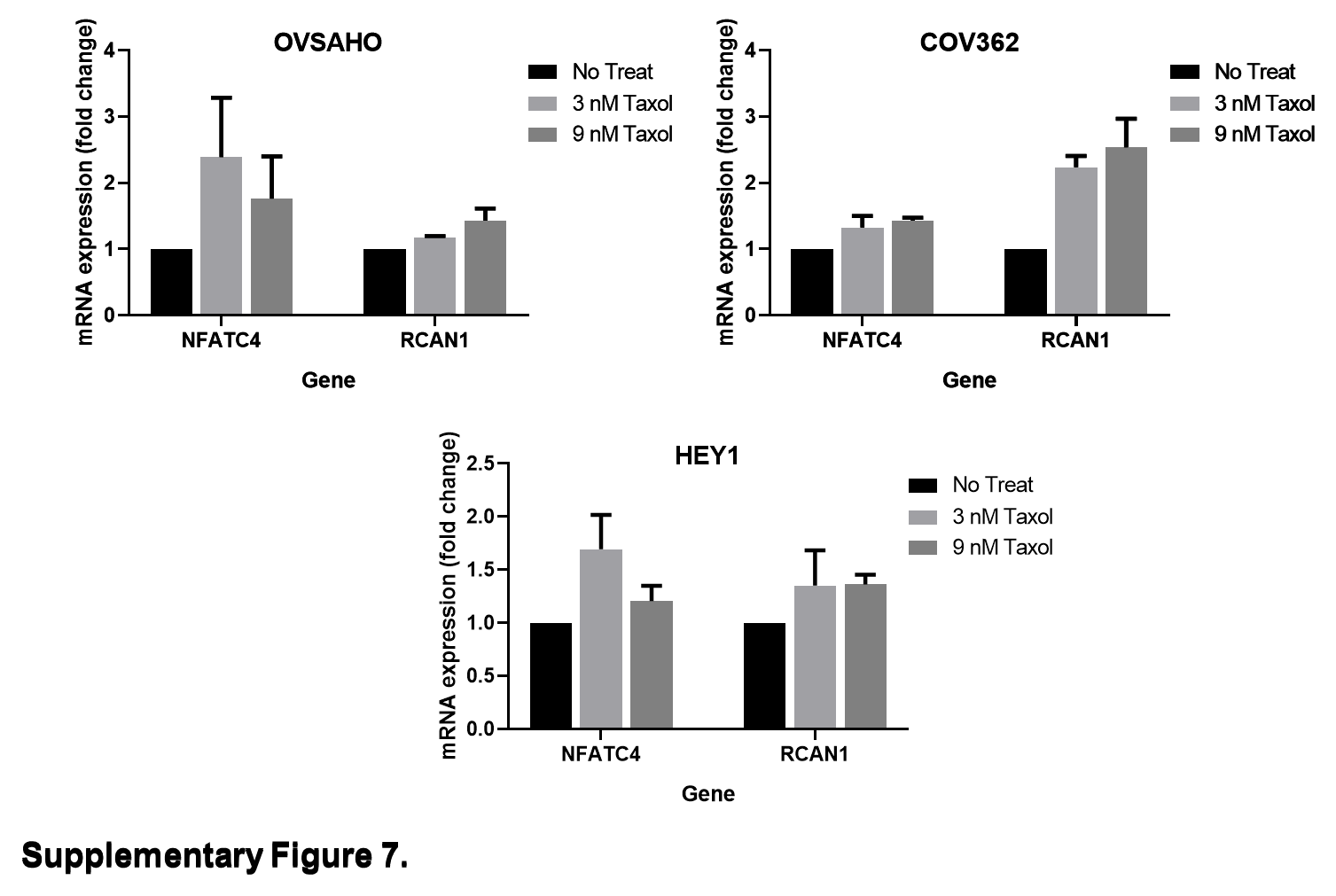

**Supplementary Figure 7**

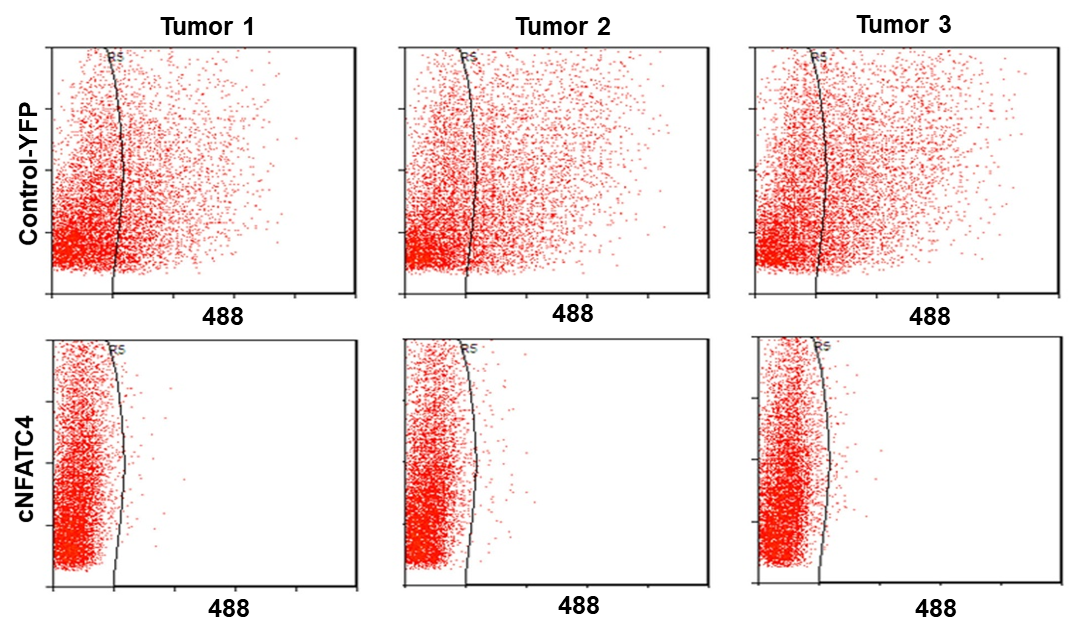

**Supplementary Figure 8**

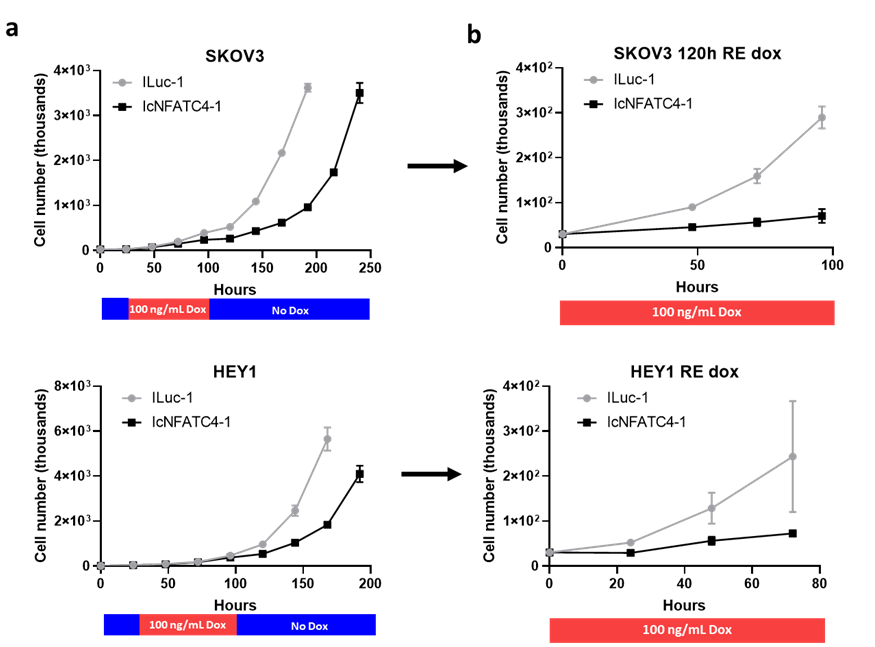

**Supplementary Figure 9**

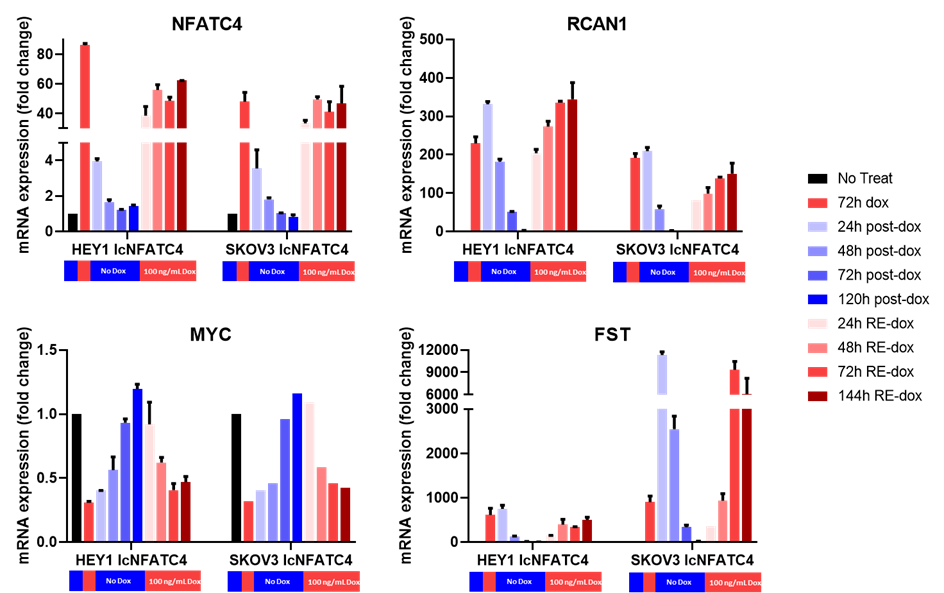

**Supplementary Figure 10**

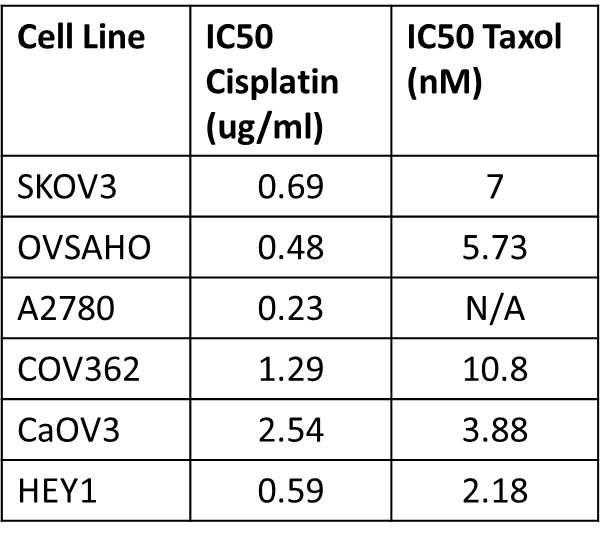

**Supplementary Table 1**

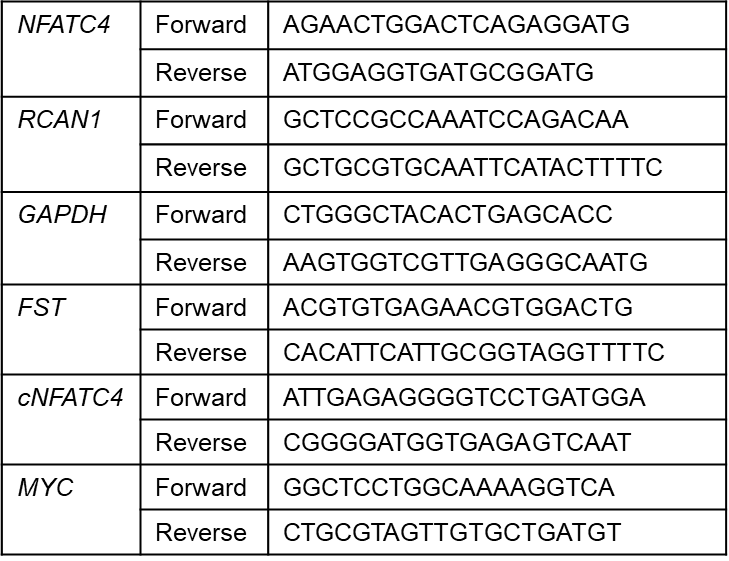

**Supplementary Table 2**
